# clootl: an R package for accessing and manipulating a complete, versioned avian phylogeny

**DOI:** 10.64898/2026.04.23.720449

**Authors:** Eliot T. Miller, Luna Luisa Sánchez Reyes, Emily Jane McTavish

## Abstract

Birds are frequently used as a focal taxon for evolutionary and ecological studies. Thousands of papers a year are published using birds as study systems. Hundreds more are published clarifying the evolutionary relationships among clades or regionally circumscribed sets of bird species. Up to date phylogenies are essential for informing and guiding avian studies, controlling for the expectation of shared trait evolution, and for science communication, among other applications. However, employing up to date phylogenies to address these questions has proven challenging for a number of reasons. First, individually published phylogenies are often hard to access in a usable manner. For example, sequences are usually made available, and an image of the phylogeny is published, but the actual phylogeny data product is often not digitally available. Second, published phylogenies often do not include all taxa of interest or have branch lengths in units of time. Third, taxonomic mismatches between phylogenies and existing datasets can complicate analyses. We address these issues by sharing an R package, ‘clootl’, that wraps together a new, complete, dated bird phylogeny with easy to use tools to extract trees for taxa of interest and sample over uncertainty. The phylogeny incorporates information from more than 300 individually published bird phylogenies. The R package includes tools to help appropriately cite these input studies. This software will enable users to smoothly integrate accurate evolutionary information into any analyses on birds.

## INTRODUCTION

The research and conservation communities rely on access to a global avian phylogeny as a key resource to provide evolutionary context for their work. Although existing examples such as Jetz et al. (2012); Burleigh et al. (2015) are widely used, these trees are static and quickly become outdated. Studies focused on smaller portions of the avian phylogeny are regularly published, revealing new insights, but these relationships need to be combined with those in other trees to be useful for large scale analyses. In addition, avian taxonomy is updated annually. These updates, if not carefully tracked, can result in a misalignment between names in databases and those in published phylogenies. We have developed a dynamic resource that can be updated and improved over time as new studies add to our collective knowledge of bird evolutionary relationships and as the taxonomy changes and species limits are redefined (McTavish et al., 2024). Here, we provide user-friendly access via an R package to this suite of linked phylogenetic and taxonomic resources.

The Open Tree of Life (OpenTree) project synthesizes published phylogenetic information to create a tree of all life. A rich suite of OpenTree software resources already exists for curating and interacting with these data, including online tools tree.opentreeoflife.org as well as R (Michonneau et al., 2016) and Python (McTavish et al., 2021) packages. While OpenTree has been successful in creating these tools and data products and making them available to the broad research community, use of these resources by the ornithology community has been limited. Dozens of bird phylogenies are published annually but, historically, few were actually curated in the OpenTree system. This lack of updated input data limited the downstream quality of the synthetic OpenTree tree, which in turn led to limited use. In addition, the flexible taxonomic approach used by OpenTree to facilitate data interoperability was a poor match to the carefully curated taxonomic treatments used by ornithologists.

Despite being static and now outdated existing resources such as Jetz et al. (2012) have had a significant and lasting impact on research and still see hundreds of published use cases per year. A dynamic, updated and high-quality phylogeny of all birds is clearly needed. To help OpenTree fulfill its potential in the avian domain and start a positive feedback loop of community buy-in, we developed a partnership between the Cornell Lab of Ornithology and OpenTree to begin careful phylogenetic and taxonomic curation of the avian phylogeny. The Cornell Lab of Ornithology (CLO) is a world-leader in the study and conservation of birds, with a mission to interpret and conserve the earth’s biological diversity through research, education, and community science focused on birds. CLO’s expertise in this domain has powered community engagement with the project. OpenTree has developed software for efficiently and reproducibly integrating large sets of phylogenetic estimates. By linking these resources together, we built and share an up-to-date, dynamic, phylogenetic estimate capturing all known bird species. This complete tree is synchronized directly with CLO’s annually updated taxonomy (Clements et al., 2025).

### THE CLOOTL PACKAGE

We built the R package ‘clootl’ to simplify access to this modern and dynamic phylogeny of the world’s birds, to facilitate research reproducibility via access to specific tree versions and taxonomies, and to ensure component datasets are properly attributed by users of the tree. This package provides access to data products from a range of taxonomic release years and phylogenetic synthesis versions. While the data products themselves are available from a GitHub repository, clootl facilitates access to the data and provides a simple interface for manipulating the larger, synthetic tree. This package meets the well-demonstrated need for a global avian phylogeny in the R environment.

Functions in the clootl package make it simple to access a dated tree for any taxon set of interest, as well as to sample across uncertainty in dates and taxonomic placements. In addition, clootl makes it straightforward to identify and properly acknowledge the constituent studies that went into creating these complete phylogenies. These citation functions can flexibly handle tree pruning, and tree and taxonomy versions, so that the acknowledged studies reflect the subset of the data output by a user. This is a key feature, since it is vital that authors of contributing studies receive credit for their work. Ensuring that constituent studies are properly attributed fosters a positive feedback loop wherein authors are incentivized to make their published phylogenies readily available for downstream use.

The package infrastructure provides a clear path to reproducibility of analyses through versioning of trees and taxonomy years. Included with the current clootl package download (v0.1.4) from CRAN or GitHub is an updated version of the bird tree of life (Aves v1.6) mapped to the new 2025 Clements taxonomy (Clements et al., 2025). This updated tree captures all 11,167 species in the 2025 Clements taxonomy (Clements et al., 2025). Of these species tips, 9,603 (86%) are placed by relationships informed by 301 published input studies. The remainder are placed using expertly curated taxonomic information. This new tree contains 38 more studies and 364 more species tips from phylogenies than the v1.4 tree published in McTavish et al. (2025). In addition, we provide R functions to download, extract and cite prior synthesis and prior taxonomy versions of the tree going back to 2021. For each taxonomy year, we provide a crosswalk table mapping species in Clements, including scientific and common names, to identifiers in the Open Tree of Life taxonomy (Rees and Cranston, 2017), and to Avibase taxon concept identifiers (Lepage et al., 2014). Where available, names are also linked to identifiers in the IOC World Bird List v14.1 (Gill et al., 2024), HBW and Birdlife v. 8.1 (HBW and International, 2023), and the Howard and Moore taxonomy (4th edition, with corrigenda from 2015) (Howard et al., 2013a,b). These taxonomy crosswalk data are included in the R package as data frames, so that relabeling tree tips from one taxonomic system to another is usually straightforward, although differences in taxon concepts across taxonomies means that a one-to-one match is not always possible. The full description of the columns in the taxonomy data files are included in the AvesData GitHub repo documentation https://github.com/McTavishLab/AvesData.

The clootl package is fully documented and well tested. Examples and tutorials demonstrate how to apply all clootl functions, and more than 75% of the code is covered by unit tests which are automatically deployed using R’s `testthat` framework (Wickham, 2011).

The package’s name is drawn from the collaboration that gave rise to it, a joint effort between the Cornell Lab of Ornithology (CLO) and the Open Tree of Life (OpenTree, or OTL) project.

### Example use cases

#### Getting a tree for an arbitrary set of species

A user can input any set of bird taxa and get a dated phylogeny for those taxa, returned as a ‘phylo’ object with branch lengths in millions of years.

~~~
ex1 <-extractTree(species=c(“Turdus migratorius”,
                            “Setophaga dominica”,
                            “Setophaga ruticilla”,
                            “Sitta canadensis”))
Alternately, users may use eBird species codes to get the tree.
ex2 <-extractTree(species=c(“amerob”,
                            “canwar”,
                            “reevir1”,
                            “yerwar”,
                            “gockin”),
                            label_type=“code”)
~~~

The columns in the taxonomy data include labels from the eBird data, such as ‘TAXON ORDER’, ‘CATEGORY’, ‘SPECIES CODE’,’TAXON CONCEPT ID’, ‘PRIMARY COM NAME’, ‘SCI NAME’, ‘ORDER’, and ‘FAMILY’. To get a tree for a higher taxon such as a family, you can pull the species names in that family from the taxonomy data and then request the tree. This example code gets a tree of all the species in the family Rhinocryptidae.

~~~
     tax <-clootl_data$taxonomies$year2025
     rhin <-tax[tax$FAMILY==“Rhinocryptidae (Tapaculos)”,]
     rhin_spp <-rhin$SCI_NAME
     ex3 <-extractTree(species=rhin_spp)
~~~

Importantly, `clootl` makes it easy to appropriately cite all studies that went into building the subtree of interest (Figure 1).

**Figure 1.**
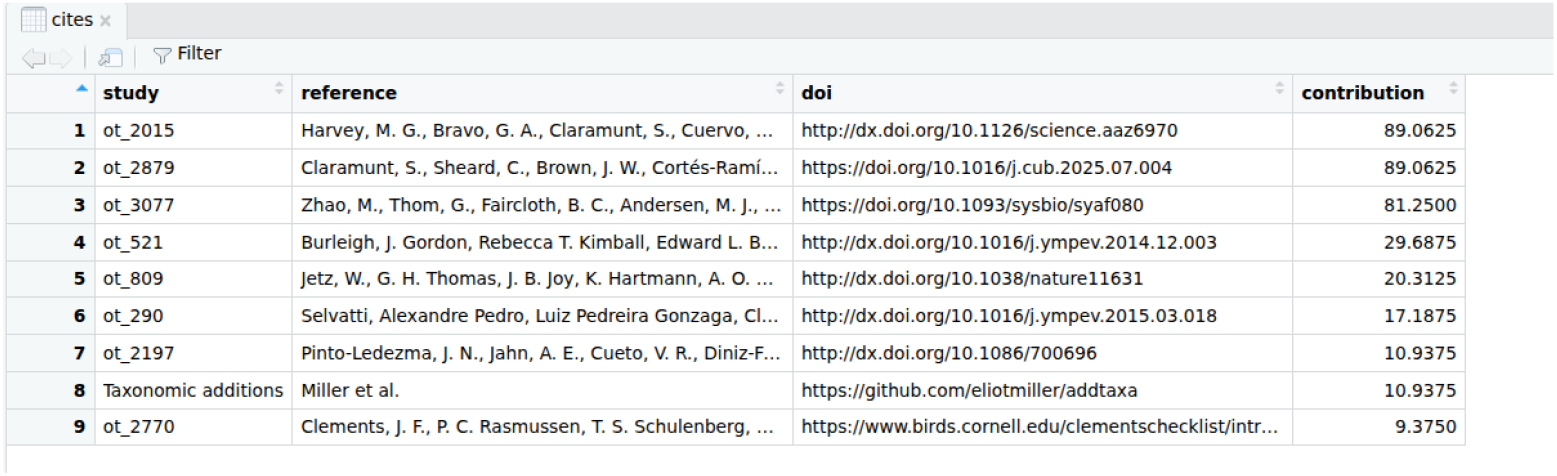
Citations for the tree of Rhinocryptidae. The weights column shows the percentage of nodes that each study contributed to relationships for.

~~~
cites<-getCitations(ex3)
~~~

The citations table returns the short form citation, digital object identifier (DOI), and the study id for each study that informed the target tree. The study id can be used to access the curated input tree on tree.opentreeoflife.org/curator. In addition, the citations information table has a ‘contribution’ column that provides a metric of each constituent study’s importance to the target tree. These values represent the percentage of nodes in the target tree informed by a given constituent study. They do not necessarily sum to 100, since multiple studies can provide information at each node. Ideally, all input studies will be cited in any downstream publication, but pragmatically this metric can be helpful when users are faced with limits on citation number. Because conflicts between studies are resolved using a greedy algorithm that ranks inputs, the fact that an input study contributed information to a node does not necessarily mean that that study agrees with the final relationships in the synthetic tree. McTavish et al. (2025) and Redelings and Holder (2017) have more detailed information on the phylogenetic synthesis procedure, and how conflict among input studies is handled. Detailed information from the OpenTree supertree synthesis including concordance and conflict between each input study and each node is packaged with the tree in the data object ‘clootl data$trees$Aves 1.6$annotations’

#### Alternate trees and taxonomies

The clootl package comes with a data object containing the most up-to-date tree version (as of this paper publication, v1.6), and the taxonomy files for all Clements taxonomies from 2021 though 2025. By default, calls to the clootl package return the most up-to-date tree and taxonomy. However, for reproducibility and for improved matches to previously published datasets (which frequently make use of older taxonomies), an optional data download is available that includes older tree versions in alternate taxonomy years. Because of the large amount of labor involved with updating taxon addition statements, over time we have deprecated older taxonomies. Therefore some taxonomies are only available for trees with older phylogenetic topology versions.

A common challenge for large scale analyses is taxononomic mismatches between datasets. We use AVONET (Tobias et al., 2022), a database of morphological measurements for the world’s birds to highlight interoperability and the ease with which clootl can be linked with existing databases. This example also serves to illustrate possible workarounds for taxonomic challenges.

To access AVONET, we download the full dataset from Figshare https://figshare.com/s/b990722d72a26b5bfead, and save the data mapped to the eBird taxonomy (sheet Avonet2 ebird) as a CSV. Using the downloaded data it is very easy to connect trait measurements to a dated phylogeny for species of interest. This example shows a sample of an arbitrary set of 50 species from AVONET.

~~~
dat <-utils::read.csv(“AVONET Supplementary dataset 1.csv”)
spp <-sample(dat$Species2, 50)
datSubset <-dat[dat$Species2 %in% spp,]
rownames(datSubset)<-datSubset$Species2
~~~

However, The AVONET dataset was published in 2021, and the data is mapped to bird species in the 2021 Clements taxonomy (Clements et al., 2019).

While the majority of species names didn’t change between 2021 and 2025, there are 589 names of the 10,661 in the AVONET data that do not match species in the 2025 taxonomy.

Users may either use the current v1.6 tree, in the 2025 taxonomy, or an older topology, such as v1.5, which is available in the 2021 taxonomy. We include a non-default ‘force’ option for users to choose drop missing taxa when matching a query species to a tree. Because the current tree will not be an exact match for the species in AVONET this ‘force’ argument is required, otherwise the call will fail due to not all query species matching.

~~~
pruned <-extractTree(species=datSubset$Species2,
                        label_type=“scientific”,
                        force=TRUE)
~~~

To extract the v1.5 tree in the 2021 taxonomy, one needs to first download the full AvesDataLite repo from GitHub using the ‘get avesdata repo’ command.

~~~
get_aves_data_repo(path=‘‘.”)
pruned <-extractTree(species=datSubset$Species2,
                        label_type=“scientific”,
                        version = 1.5,
                        taxonomy_year=2021)
~~~

Using either of these trees matched to the AVONET data we can easily analyze and plot trait changes through evolutionary time. Here we show a continuous stochastic character map of the natural logarithm of body mass (Figure 2).

**Figure 2.**
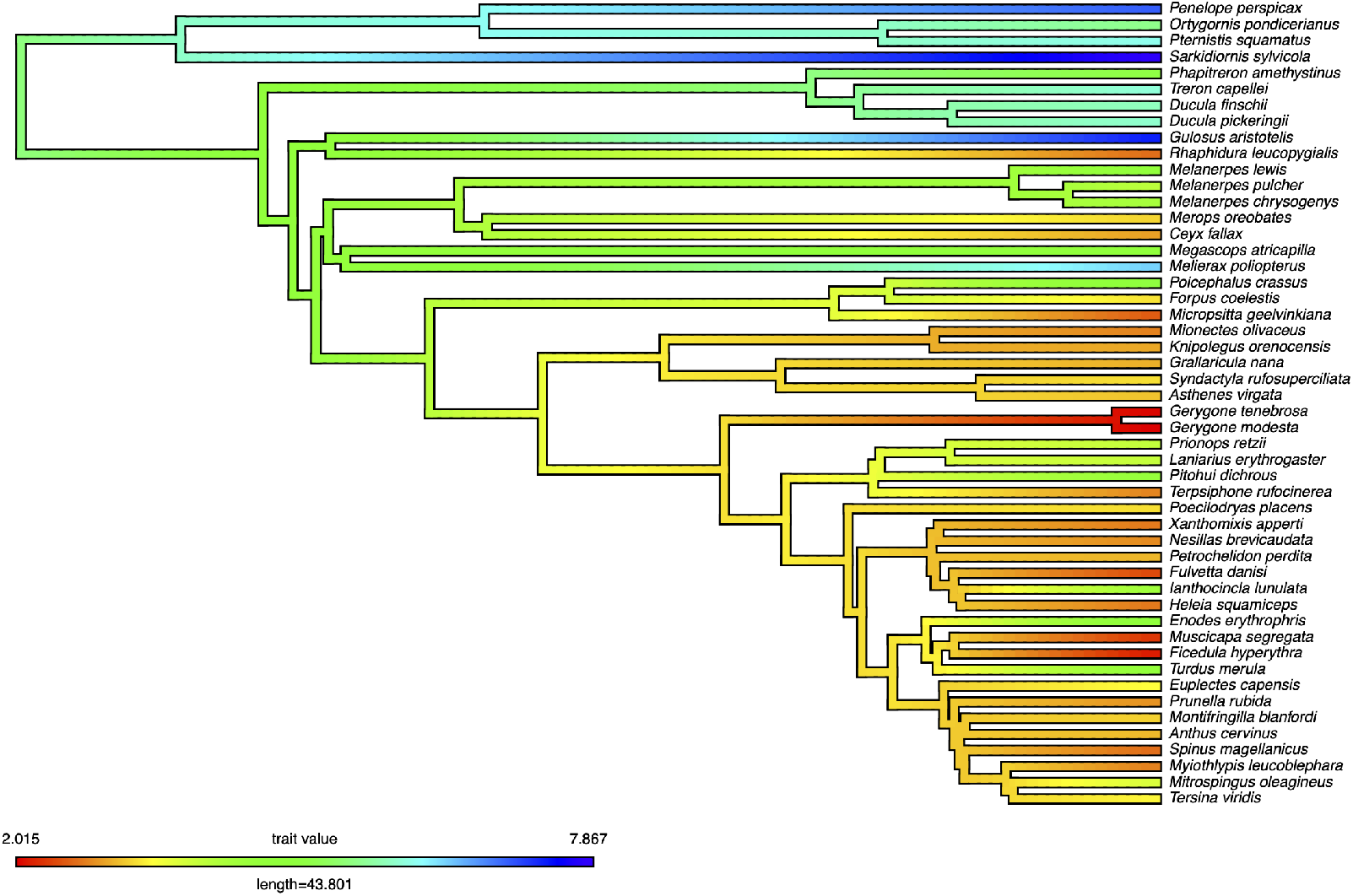
A continuous stochastic character map of the natural logarithm of body mass for 50 randomly sampled bird species. This example used the v1.5 tree in the 2021 taxonomy for an exact match to the AVONET data

#### Sampling over trees

The trees used in clootl are generated following the methods described in McTavish et al. (2024). While by default clootl functions reference the single summary tree for each phylogenetic synthesis and taxonomy year, we have also built functionality to capture topological and divergence date uncertainty in our outputs.

This uncertainty is intentionally incorporated into two parts of the tree-building process. The first is the taxonomic addition step, when the approximately 15% of taxa that are not present in any of the input phylogenies are added stochastically to the backbone tree based on placement constraints developed using expert knowledge (Mast et al., 2015). This stochastic placement step is repeated 100 times, generating 100 trees in which the core phylogenetic backbone relationships remain the same, but these taxon additions vary. Then, each of these trees are individually dated by resampling from the distributions of input dates in our tree store for each node using Chronosynth (McTavish and Sánchez Reyes, 2026; Sánchez Reyes et al., 2024). This date resampling process captures some of the uncertainty in node ages across the tree.

The function ‘sampleTrees()’ in clootl makes it easy to get a sample of 100 trees capturing this stochastic variation in relationships and node ages for any taxon set of interest.

~~~
sampleA <-sampleTrees(species=c(“Turdus migratorius”,
                                        “Setophaga dominica”,
                                        “Setophaga ruticilla”,
                                        “Sitta canadensis”))
## Here there is only variation in node ages phangorn::densiTree(sampleA)
## For the genus Mergus some species were placed taxonomically ## This results in variation in ages and relationships
tax <-clootl_data$taxonomies$year2025
genus<-stringr::str_split_i(tax$SCI_NAME, “ “,1)
tax<-cbind(tax, GENUS=genus)
Mergus <-tax[tax$GENUS==“Mergus”,]
Mergus_spp <-Mergus$SCI_NAME
sampleB <-sampleTrees(species=Mergus_spp)
phangorn::densiTree(sampleB)
~~~

For sets of taxa which are entirely included in input phylogenies, the variation across sampled trees will only be in branch length (e.g. sampleA in the example code, Figure 3a), whereas for other groups where more taxa have been added based on taxonomic information, there is variation in relationships and in ages (sampleB in the example code, Figure 3b). In the second example for the five species in the genus Mergus, *Mergus australis* (Aukland Islands Merganser) and *Mergus octosetaceus* (Brazilian merganser) are not included in any input phylogenies, and as such are placed stochastically within the genus. By iteratively re-running analyses over a sample of trees capturing this topological and dating uncertainty, the robustness of conclusions can be assessed.

**Figure 3.**
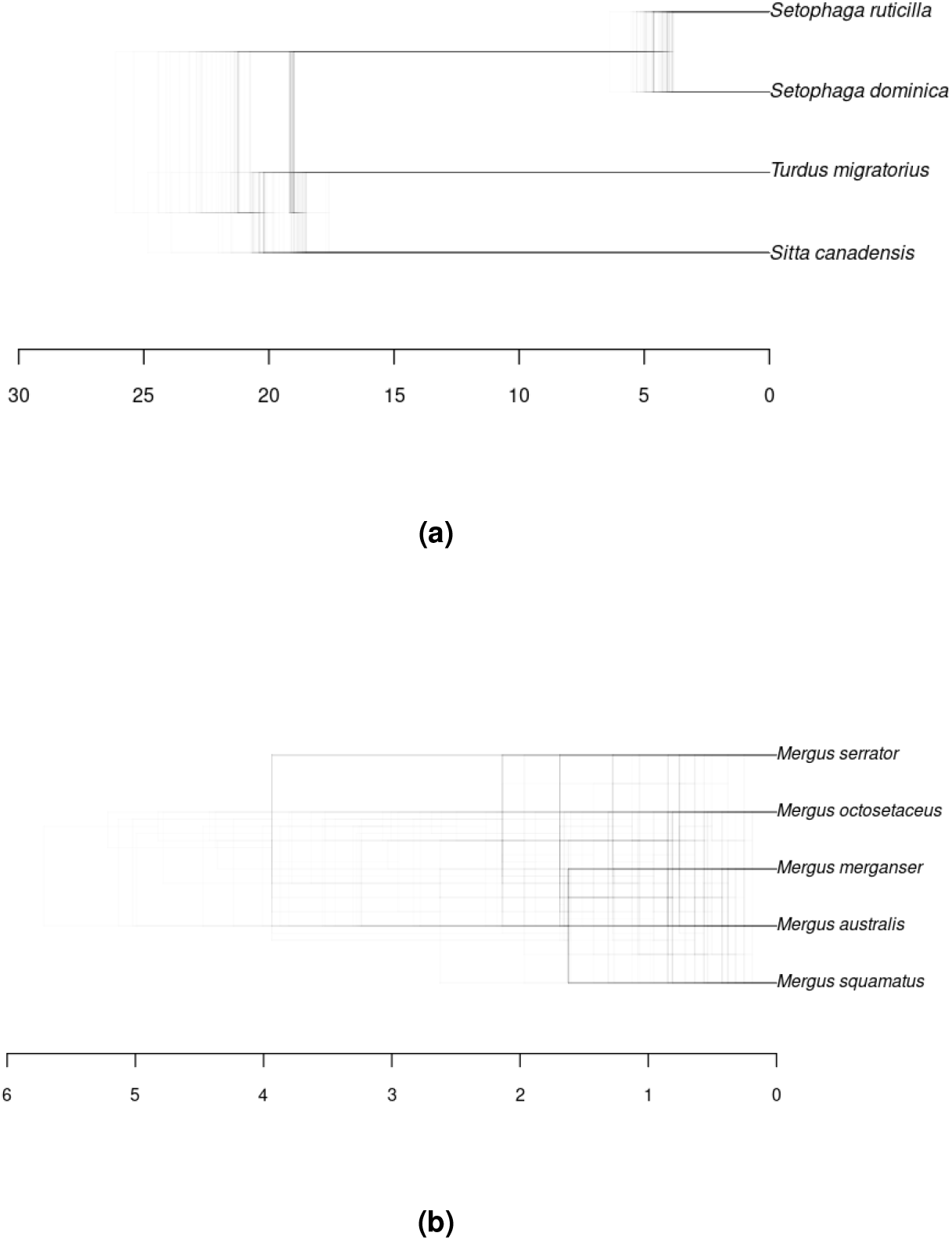
A) Denstiree plot showing variation in branch length across 100 sampled trees for four species. Code shown B) Denstiree plot showing variation in topology and branch length across 100 sampled trees for 5 species in the genus Mergus. *Mergus australis*(Aukland Islands Merganser) and *Mergus octosetaceus* (Brazilian merganser) are not sampled in any input phylogenies, and are placed stochastically based on taxonomic information. Branch lengths are in millions of years. Figure made using phangorn (Schliep, 2011)

## CONCLUSION

Our package, clootl, provides user friendly access to a complete, updated tree of all birds, including taxonomy files, synthetic phylogenies in multiple versions, sets of trees capturing uncertainty in estimates and in dates, and associated bibliographic information about the inputs. clootl offers direct and easy access to a new and perpetually improving phylogeny of the world’s birds.

## Availability and Reproducibility

Software is available from CRAN and from Github at https://github.com/eliotmiller/clootl. Code to reproduce analyses in this paper is available in the clootl GitHub repo. All datasets are posted on GitHub.

## ACKNOWLEDGMENTS

EJM was supported by NSF ABI #1759846. We thank Jimmy Choi and Robin Trayler for assistance with testing.

## AUTHOR CONTRIBUTION STATEMENT

All three authors contributed to the code development, documentation, testing and manuscript writing.

---

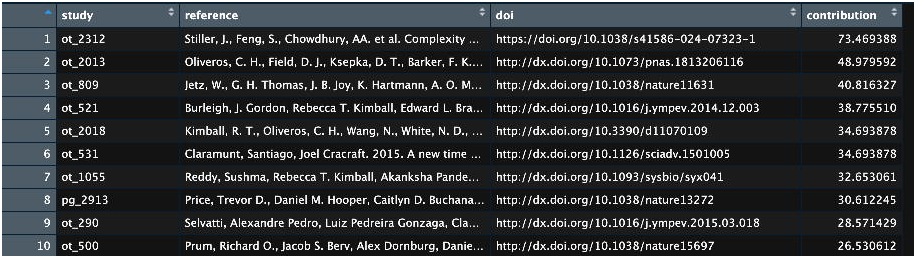

## Notes

https://github.com/eliotmiller/clootl

### Competing Interest Statement

The authors have declared no competing interest.

## REFERENCES

Burleigh, J. G., Kimball, R. T., and Braun, E. L. (2015). Building the avian tree of life using a large-scale, sparse supermatrix. Molecular Phylogenetics and Evolution, 84:53–63.

Clements, J. F., Rasmussen, P. C., Schulenberg, T. S., Iliff, M. J., Gerbracht, J. A., Lepage, D., Spencer, A., Billerman, S. M., Sullivan, B. L., Smith, M., and Wood, C. L. (2025). The ebird/clements checklist of birds of the world: v2025. Downloaded from https://www.birds.cornell.edu/clementschecklist/download/.

Clements, J. F., Schulenberg, T. S., Iliff, M. J., Billerman, S. M., Fredericks, T., Sullivan, B. L., and Wood, C. L. (2019). The eBird/Clements checklist of birds of the world: v2019.

Gill, F., Donsker, D., and Rasmussen, P. (2024). Ioc world bird list (v14. 1). 10.14344/ioc.

HBW and International, B. (2023). Handbook of the birds of the world and birdlife international digital checklist of the birds of the world. version 8.

Howard, R., Moore, A., Dickinson, E. C., Remsen, J. V., Cracraft, J., and Christidis, L. (2013a). The Howard and Moore Complete Checklist of the Birds of the World: Passerines. Aves Press, United Kingdom.

Howard, R., Moore, A., Dickinson, E. C., Remsen, J. V., Cracraft, J., and Christidis, L. (2013b). The Howard and Moore Complete Checklist of the Birds of the World: Passerines. Aves Press, United Kingdom.

Jetz, W., Thomas, G. H., Joy, J. B., Hartmann, K., and Mooers, A. O. (2012). The global diversity of birds in space and time. Nature, 491(7424):444–448.

Lepage, D., Vaidya, G., and Guralnick, R. (2014). Avibase – a database system for managing and organizing taxonomic concepts. ZooKeys, (420):117–135.

Mast, A. R., Olde, P. M., Makinson, R. O., Jones, E., Kubes, A., Miller, E. T., and Weston, P. H. (2015). Paraphyly changes understanding of timing and tempo of diversification in subtribe hakeinae (proteaceae), a giant australian plant radiation. American Journal of Botany, 102(10):1634–1646.

McTavish, E. J., Gerbracht, J. A., Holder, M. T., Iliff, M. J., Lepage, D., Rasmussen, P., Redelings, B., Reyes, L. L. S., and Miller, E. T. (2024). A complete and dynamic tree of birds. Pages: 2024.05.20.595017 Section: New Results.

McTavish, E. J., Gerbracht, J. A., Holder, M. T., Iliff, M. J., Lepage, D., Rasmussen, P. C., Redelings, B. D., Sánchez Reyes, L. L., and Miller, E. T. (2025). A complete and dynamic tree of birds. Proceedings of the National Academy of Sciences, 122(18):e2409658122. Publisher: Proceedings of the National Academy of Sciences.

McTavish, E. J. and Sánchez Reyes, L. L. (2026). Chronosynth. https://github.com/OpenTreeOfLife/chronosynth.

McTavish, E. J., Sánchez-Reyes, L. L., and Holder, M. T. (2021). Opentree: A python package for accessing and analyzing data from the open tree of life. Systematic Biology, 70(6):1295–1301.

Michonneau, F., Brown, J. W., and Winter, D. J. (2016). rotl: an r package to interact with the open tree of life data. Methods in Ecology and Evolution, 7(12):1476–1481.

Redelings, B. D. and Holder, M. T. (2017). A supertree pipeline for summarizing phylogenetic and taxonomic information for millions of species. PeerJ, 5:e3058.

Rees, J. A. and Cranston, K. (2017). Automated assembly of a reference taxonomy for phylogenetic data synthesis. Biodiversity Data Journal.

Sánchez Reyes, L. L., McTavish, E. J., and O’Meara, B. (2024). Datelife: leveraging databases and analytical tools to reveal the dated tree of life. Systematic Biology, 73(2):470–485.

Schliep, K. P. (2011). phangorn: phylogenetic analysis in R. Bioinformatics, 27(4):592–593.

Tobias, J. A., Sheard, C., Pigot, A. L., Devenish, A. J. M., Yang, J., Sayol, F., Neate-Clegg, M. H. C., Alioravainen, N., Weeks, T. L., Barber, R. A., Walkden, P. A., MacGregor, H. E. A., Jones, S. E. I., Vincent, C., Phillips, A. G., Marples, N. M., Montaño-Centellas, F. A., Leandro-Silva, V., Claramunt, S., Darski, B., Freeman, B. G., Bregman, T. P., Cooney, C. R., Hughes, E. C., Capp, E. J. R., Varley, Z. K., Friedman, N. R., Korntheuer, H., Corrales-Vargas, A., Trisos, C. H., Weeks, B. C., Hanz, D. M., Töpfer, T., Bravo, G. A., Remeš, V., Nowak, L., Carneiro, L. S., Moncada R., A. J., Matysioková, B., Baldassarre, D. T., Martínez-Salinas, A., Wolfe, J. D., Chapman, P. M., Daly, B. G., Sorensen, M. C., Neu, A., Ford, M. A., Mayhew, R. J., Fabio Silveira, L., Kelly, D. J., Annorbah, N. N. D., Pollock, H. S., Grabowska-Zhang, A. M., McEntee, J. P., Carlos T. Gonzalez, J., Meneses, C. G., Muñoz, M. C., Powell, L. L., Jamie, G. A., Matthews, T. J., Johnson, O., Brito, G. R. R., Zyskowski, K., Crates, R., Harvey, M. G., Jurado Zevallos, M., Hosner, P. A., Bradfer-Lawrence, T., Maley, J. M., Stiles, F. G., Lima, H. S., Provost, K. L., Chibesa, M., Mashao, M., Howard, J. T., Mlamba, E., Chua, M. A. H., Li, B., Gómez, M. I., Garcíía, N. C., Päckert, M., Fuchs, J., Ali, J. R., Derryberry, E. P., Carlson, M. L., Urriza, R. C., Brzeski, K. E., Prawiradilaga, D. M., Rayner, M. J., Miller, E. T., Bowie, R. C. K., Lafontaine, R.-M., Scofield, R. P., Lou, Y., Somarathna, L., Lepage, D., Illif, M., Neuschulz, E. L., Templin, M., Dehling, D. M., Cooper, J. C., Pauwels, O. S. G., Analuddin, K., Fjeldså, J., Seddon, N., Sweet, P. R., DeClerck, F. A. J., Naka, L. N., Brawn, J. D., Aleixo, A., Böhning-Gaese, K., Rahbek, C., Fritz, S. A., Thomas, G. H., and Schleuning, M. (2022). AVONET: morphological, ecological and geographical data for all birds. Ecology Letters, 25(3):581–597. eprint: https://onlinelibrary.wiley.com/doi/pdf/10.1111/ele.13898.

Wickham, H. (2011). testthat: Get started with testing. The R Journal, 3:5–10.

